# DDX5 and DDX17 RNA helicases regulate hepatitis B virus RNA splicing

**DOI:** 10.1101/2025.02.14.638243

**Authors:** Guillaume Giraud, Xavier Grand, Pélagie Huchon, Mélanie Rodà, Audrey Diederichs, Fleur Chapus, Francesca De Nicola, Romain Parent, Michel Rivoire, Cyril F. Bourgeois, Fabien Zoulim, Barbara Testoni

## Abstract

Chronic HBV infection remains a major health burden worldwide and is the main driver of severe liver diseases. Liver pathogenesis is associated with the increased proportion of HBV spliced variants that encode viral proteins involved in liver disease progression. However, how HBV RNA splicing is regulated is poorly understood. Here, we focused on DDX5 and DDX17 RNA helicases, known to regulate HBV RNA metabolism and alternative splicing of host genes. By performing 5’RACE-PCR combined with single molecule sequencing, we demonstrated that silencing both proteins increased the usage of a specific splicing donor site and the expression of the derived HBV spliced variants. Polysome fractionation highlighted the ability of these RNA species to encode new viral proteins potentially contributing to liver pathogenesis. Overall, our data established DDX5 and DDX17 helicases as master regulators of HBV RNA metabolism, by fine-tuning viral splicing, which is linked to HBV-induced liver pathogenesis and disease progression.

## INTRODUCTION

Despite an efficient prophylactic vaccine, chronic hepatitis B virus (HBV) infection remains a major health burden worldwide. CHB patients are at high risk of developing severe liver diseases such as fibrosis, cirrhosis, and eventually hepatocellular carcinoma (HCC) ^1^. Current standard-of-care treatment based on nucleos(t)ide analogs or pegylated interferon-α are efficient in maintaining the infection under control and lowering the risk of liver disease, however chronic HBV carriers still present a higher risk to develop HCC respect to the non-infected individuals. Furthermore, these treatments cannot completely eradicate the virus due to the persistence of its minichromosome, the so-called covalently closed circular (ccc)DNA ^1^. This chromatinised episome is the template for the transcription by the host RNA polymerase II (RNAP II) of the six main viral mRNAs, including the pregenomic (pg)RNA that is retro-transcribed by the viral polymerase to generate new infectious viral particles ^2–4^.

As host RNAP II-synthesised transcripts, pgRNA and the PreS2/S RNA, encoding the envelope proteins, can undergo alternative splicing. Up to date, twenty-two spliced variants (SPs) have been identified in patient’s serum and in *in vitro* cell lines. Each of the twenty SPs (SP01 to SP20) derived from pgRNA and the two (SP21 and SP22) derived from PreS2/S RNAs differ by the specific combination of splicing junctions between donor and acceptor sites. However, a single splicing junction can be found in several SPs making their particular detection and quantification difficult by classical molecular biology approaches ^5^. pgRNA-derived SPs are retro-transcribed and generate new viral particles that cannot initiate a new replication cycle ^6^. As a consequence, HBV RNA splicing is considered non-essential for viral replication, contrary to other viruses such as HIV-1 ^7^. Nonetheless, recent studies highlighted the importance of HBV RNA splicing-derived proteins in fine-tuning HBV replication ^5,8–11^. Interestingly, the proportion of SPs in the blood of CHB patients appears to increase one to three years prior to the development of HCC ^12^. This observation suggests that, next to their role in viral replication, SPs also play a role in the liver disease induced by HBV. Recently, *in vitro* co-culture assays between hepatic stellate cells (HSC) and hepatocytes expressing HBV SPs (SP01 to SP05) showed that SPs can activate HSCs, which are known to induce liver fibrosis ^13^. Again, SP-encoded proteins are thought to mediate the contribution of the SPs to liver pathogenesis. The SP01-derived HBSP (Hepatitis B Spliced Protein) protein was demonstrated to inhibit the host immune response against HBV and to activate pathways involved in carcinogenesis ^14–16^. Besides, HBSP was associated with HCC metastasis and increased cell motility most likely by interacting with the lysosomal protease, cathepsin B ^17^. Furthermore, HBV SPs are considered as an additional source of the HBx protein that is a key driver of HCC ^18,19^. Given the pleiotropic role that SPs play on both HBV replication and induced liver disease, identifying the host factors regulating HBV RNA splicing is thus fundamental in the aim of designing strategies to modulate their expression and potentially influence patients’ outcomes. So far, few host factors, such as TARDBP or PSF have been identified regulating this important process ^5,10,20,21^.

DDX5 and DDX17 RNA helicases have redundant functions in controlling almost all stages of gene expression ^22,23^. In particular, both proteins regulate alternative splicing of host cell transcripts by controlling the recognition of splicing donor sites by the spliceosome ^22,24–28^. Next to their role in host gene expression, DDX5 and DDX17 appear to be either proviral or antiviral factors for several viruses by regulating viral gene expression, including splicing of viral transcripts ^27,29,30^. Accordingly, we and others pointed out the role of DDX5 and DDX17 as host restriction factor for HBV ^31–33^. In particular, we recently demonstrated that, among the different mechanisms orchestrated by both helicases to inhibit HBV replication, DDX5 and DDX17 directly regulate the co-transcriptional 3’ end HBV RNA processing ^31^, establishing a clear role of both proteins as key regulators of HBV RNAs metabolism. Interestingly, the expression of both helicases is down-regulated in CHB and HBV-induced HCC patients and lower expression of DDX5 is associated with a poor prognosis ^31,33^.

Here, we investigated the role of DDX5 and DDX17 in HBV RNA splicing. By combining 5’RACE-PCR and Oxford Nanopore Technologies-based sequencing of RNAs purified from DDX5 and DDX17-silenced HBV-infected HepG2-NTCP cells, we demonstrated that both helicases prevent the recognition of the splicing donor site at position 2070. Consequently, silencing of both proteins is associated with the increased production of SP02, SP03, SP15, SP16 and SP17. Furthermore, we performed polysome fractionation and demonstrated that these SPs are translated in HepG2-NTCP cells indicating that additional HBV proteins potentially involved in liver pathogenesis can be produced by SPs.

## RESULTS

### DDX5 and DDX17 silencing increases the production of specific HBV SPs

To unveil the role of DDX5 and DDX17 RNA helicases in HBV RNA splicing, we analysed the HBV transcriptome by performing 5’RACE-PCR assays combined with single molecule sequencing in DDX5 and DDX17 (DDX5/17) silenced HepG2-NTCP cells infected with HBV for 8 days (**Figure S1a**). We previously demonstrated that this method is able to discriminate individual HBV transcripts ^4^. We then sequenced 5’RACE amplicons (**Figure S1b**) at the single-molecule level using the Oxford Nanopore Technology (ONT). Sequencing data were analysed using our recently developed Bolero pipeline that is able to quantify the proportion of each individual HBV RNAs, including HBV SPs, related to overall number of reads aligning to the HBV genome ^34^. We first determined whether DDX5 and DDX17 influence the proportion of SPs in HepG2-NTCP cells. Each SP can be discriminated from each other by their transcription start site (TSS) position and their unique combination of splicing junction. As an example, while SP01 and SP02 share the same TSS and the splicing junction J2449-491, they differ by the splicing junction J2069-2352, present in SP02 (**Figure 1a**). Thanks to these specific features, Bolero can discriminate and compare the proportion of each individual SPs across different conditions. Interestingly, DDX5/17 silencing was associated with a significant increase in the proportion of the pgRNA-derived SP03 and SP17 (**Figure 1b, S1c**). Strikingly, the proportion of SP17, which is very low in control cells (0.37 %), strongly increased to 11.25% in silenced cells. The decrease in the proportion of other SPs in DDX5/17 silenced cells, despite not reaching statistical significance, could account for a compensation leading to the absence of difference in the global proportion of SPs between control and DDX5-17 silenced cells (**Figure 1c, S1c**). These results suggest then that DDX5 and DDX17 regulate HBV RNA splicing qualitatively rather than quantitatively.

**Figure 1:**
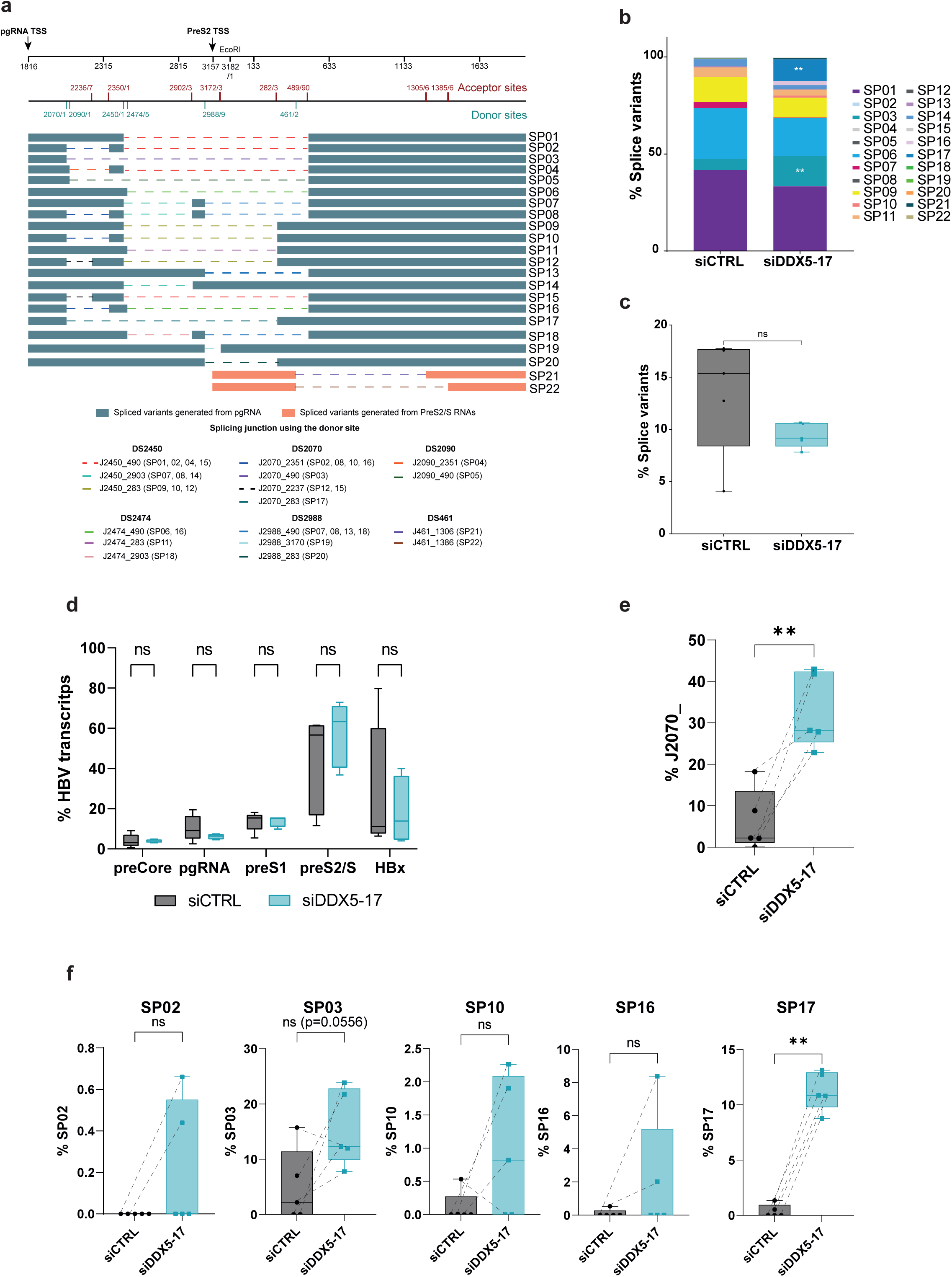
DDX5 and DDX17 silencing regulate HBV RNA splicing by preventing the recognition of the donor site 2070. **a** Scheme of the different spliced variants derived from pgRNA (green) or PreS2/S RNAs (orange) (Adapted from Grand *et al.*, 2024). The +1 position corresponds to the EcoRI cut site. The coordinates correspond to those of the HBV genotype D substrain ayw. The TSS of pgRNA and PreS2/S RNAs are indicated by an arrow. Each splicing junctions are in dashed lines. **b-f** HepG2-NTCP cells were infected with HBV and transfected twice with control siRNAs (siCTRL) or siRNAs directed against DDX5 and DDX17 mRNAs (siDDX5-17). 8 days post-infection, RNAs were isolated and subjected to 5’ RACE-PCR. **b** Proportion of each HBV SP identified thanks to their splicing junction and their TSS normalised to their original transcript (pgRNA for SP1 to SP20 SPs and PreS2/S for SP21 and SP22). **c** Percentage of the number of reads aligned to HBV containing a splicing junction normalised to the total number of reads aligned to HBV in siCTRL (grey boxplot) and in siDDX5-17 conditions (green boxplot). Dashed lines connect each biological replicate. **c** Percentage of the different HBV transcripts based on the TSS usage count normalised on the total number of reads aligned to HBV. **d** Percentage of the HBV reads containing the SD2070 engaged in a splicing junction over the total number of pgRNA reads in siCTRL (grey boxplot) and siDDX5-17 conditions (green boxplot). Dashed lines connect each biological replicate. **e** Percentage of SP02, SP03, SP10, SP16 and SP17 SPs containing the SD2070 engaged in a splicing junction over the total number of pgRNA reads in siCTRL (grey boxplot) and siDDX5-17 conditions (green boxplot). Dashed lines connect each biological replicate. **c-f** Boxplots represent the minimum, the median, and the maximum values of five independent biological replicates. Kruskal Wallis (panel **b**) and Mann-Whitney (panel **a**, **c-f**) were performed to compare siCTRL and siDDX5-17 conditions. ns: p>0.05; *: p<0.05; **: p<0.01.

In parallel, we also looked at the proportion of the main viral mRNAs that can be discriminated by their unique TSS. Both PreS2 and S TSS were considered as unique due to their close proximity sequence-wise, as was the case for HBx TSSs ^34^. As observed in **Figure S1d**, the TSS usage profiles of both control and DDX5/17 silenced cells were very similar. Consistently, the quantification of the individual viral transcripts also revealed no significant changes, indicating that DDX5 and DDX17 did not influence the TSS usage of the different HBV transcripts (**Figure 1d**). Taken together, these data showed that DDX5 and DDX17 regulate the production of specific SPs in infected HepG2-NTCP cells.

### DDX5 and DDX17 silencing is associated with the differential usage of splicing donor sites

These results pointed out the specific increased proportion of SP03 and SP17 upon DDX5/17 silencing in HepG2-NTCP cells. Interestingly, both SP03 and SP17 are characterized by splicing junctions involving the splicing donor site at the position 2070 (DS2070) (**Figure 1a**) suggesting that DDX5 and DDX17 might prevent the recognition of this specific site. Accordingly, the quantification of Nanopore reads containing DS2070 engaged in a splicing junction revealed a significant increased usage of this specific site upon DDX5/17 silencing compared to control cells (**Figure 1e**), differently from other pgRNA donor sites (DS2450, DS2474, DS2988 and DS2090) (**Figure S1e**). In addition to SP03 and SP17, the DS2070 is also engaged in splicing junction generating the SP02, SP08, SP10, SP12, SP15 and SP16. SP08, SP12, and SP15 were never detected in our experimental conditions, while SP02, SP10, and SP16 were expressed at very low levels and tended to increase following DDX5/17 silencing (**Figure 1f, S1c**). Overall, these data suggest that DDX5 and DDX17 prevent the recognition of the DS2070.

We also noticed the reproducible increase in the usage of the PreS2/S DS461 upon DDX5/17 silencing (**Figure S1e-f**). This DS is engaged in splicing junctions generating 2 PreS2/S-derived SPs, SP21 and SP22 that are scantly expressed in HepG2-NTCP cells ^5,34^. Accordingly, SP21 and SP22 proportion tended to increase in DDX5/17 silenced cells, suggesting that both helicases also prevent the recognition of this DS (**Figure S1f**). Taken together, these data demonstrate that DDX5 and DDX17 regulate HBV RNA splicing by potentially preventing the usage of specific splicing donor sites.

### Validation of the role of DDX5 and DDX17 in preventing the recognition of DS2070

We then sought to validate the regulation of the SD2070 usage by DDX5 and DDX17 in HepG2-NTCP cells by two alternative approaches. The first approach consisted in analysing RNA-seq data obtained from HepG2-NTCP cells silenced or not for DDX5 and DDX17. As we did with our 5’RACE approach (**Figure 1e**), we quantified the number of reads containing the DS2070 engaged in a splicing junction and normalised them to the total number of reads aligning to the HBV genome. As previously observed, silencing of DDX5 and DDX17 is associated with the significant increased usage of the DS2070 compared to control cells (**Figure 2a**) while the usage of the other splicing sites was not significantly modified in the same condition (**Figure S2**).

**Figure 2:**
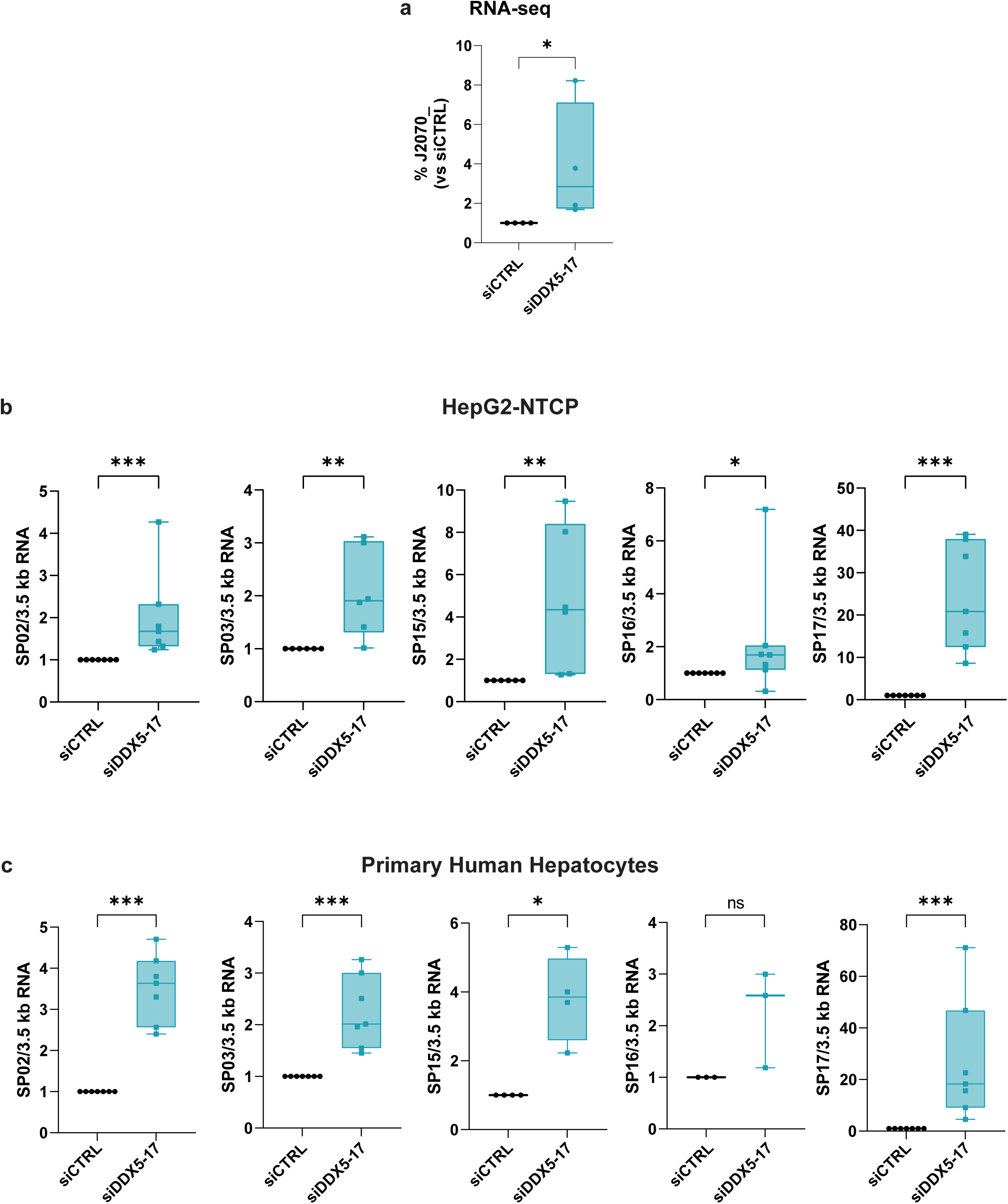
Validation of the 5’RACE PCR assays. **a-b** HepG2-NTCP cells were treated as in Figure 1. **a** GSE239571 RNA-seq dataset was reanalysed completed with two other biological replicates. Reads containing the SD2070 engaged in a splicing junction were counted and normalised to the total number of reads aligning to HBV in siCTRL (grey boxplot) and siDDX5-17 (green boxplot) conditions. Dashed lines connect each biological replicate. **b** cDNAs were subjected to ddPCR using junction primers specific to SP02, SP03, SP15, SP16 and SP17 HBV SPs. Signals were normalised to the signals obtained by ddPCR amplification of the 3.5 kb RNAs and to the siCTRL condition. **c** RNAs isolated from PHH obtained from three to four different donors treated as HepG2-NTCP cells were subjected to RT-ddPCR as in **b**. **a-c** Boxplots represent the minimum, the median, and the maximum values of four (**a**) six to seven (**b**) and three to four (**c**) independent biological replicates. Mann-Whitney tests were performed to compare siCTRL and siDDX5-17 conditions. ns: p>0.05; *: p<0.05; **: p<0.01; ***: p<0.005.

The second approach consisted in quantifying specifically each individual SPs by digital droplet PCR (ddPCR). To this purpose, we designed specific junction primers that span splicing junctions (**Figure S3a**). We checked first the specificity of these primers by performing ddPCR from HepG2-NTCP cells infected or not (MOCK control) with HBV. As observed in **Figure S3b**, signals derived from amplification by the primers specific to SP02, SP03, SP16 and SP17 were only detected in the HBV condition, as those derived from the primers used to quantify the 3.5 kb RNAs (used as signal normaliser). Thanks to the high sensitivity of the ddPCR approach, we were able to detect a specific amplification signal for SP15 in the HBV condition (**Figure S3b**). A similar approach was used to quantify the SP10 variant but we detected unspecific signal in the MOCK control (data not shown). Thus, we decided to quantify SP02, SP03, SP15, SP16 and SP17 in HepG2-NTCP cells infected and silenced for both DDX5 and DDX17 by using our ddPCR specific approach. Strikingly, silencing of both RNA helicases was associated with the significant increase of the levels of the above-mentioned SPs compared to control cells (**Figure 2b**). As already observed by 5’RACE (**Figure 1b, S1c**), SP17 was the SP that displayed the strongest induction with a fold change of approximately 20-fold in average in DDX5/17 silenced cells compared to control cells (**Figure 2b**). These data are thus in agreement with our previous 5’RACE and RNA-seq analyses showing the increased proportion of the SPs carrying spliced junctions involving the DS2070.

HBV RNA splicing was shown to be modulated depending on the cell type in which HBV replicates ^35^. Furthermore, cancer cells are associated with alternative splicing alterations due to the differential action of splicing factors ^36^. Thus, we sought to determine whether the contribution of DDX5 and DDX17 to HBV RNA splicing was conserved in primary human hepatocytes (PHH). We thus infected PHH isolated from different donors with HBV for 8 days and silenced both DDX5 and DDX17 expression. We then performed ddPCR to specifically quantify the levels of SP02, SP03, SP15, SP16 and SP17. Interestingly, as observed in HepG2-NTCP cells, the levels of all these SPs were significantly increased in DDX5/17-silenced PHH compared to control cells (**Figure 2c**). Again, SP17 was the SP showing the highest induction with a fold change of 20 between silenced and control PHH (**Figure 2c**). These data thus demonstrate that the contribution of DDX5 and DDX17 to HBV RNA splicing is conserved in PHH, the gold standard for studying HBV natural infection.

Overall, these results strongly suggest that DDX5 and DDX17 prevent the recognition of the DS2070 in HepG2-NTCP cells and PHH. Consequently, DDX5 and DDX17 silencing results in significantly increased proportion of SPs carrying a spliced junction involving the DS2070, SP17 being the mostly induced SPs.

### SPs containing the DS2070 potentially encode new HBV proteins

Some HBV SPs have been shown to encode new HBV proteins that not only contribute to viral replication but also to HBV-induced liver pathogenesis ^5,10^. We thus wondered whether the SPs regulated by DDX5 and DDX17 might also encode new viral proteins. To answer this question, we performed polysome fractionation from HepG2-NTCP cells infected and silenced for both helicases and quantified by ddPCR the levels of SP02, SP03, SP15, SP16 and SP17 associated to the non-polysomal and polysomal fractions, the latter containing translated mRNAs. Before loading the lysate to the sucrose gradient, part of the samples was treated with EDTA that releases mRNAs from ribosomes, hence providing a biochemical proof for genuine ribosomal origin of the quantified signals ^37^. Furthermore, the lack of a 18S signal in the EDTA negative control indicate that EDTA efficiently released functional ribosomes from mRNAs. We then submitted the RNAs purified from each fractions to an RNA electrophoresis in order to locate polysomal fractions, characterized by a 28S/18S rRNAs ratio of 1.7 that correspond to the fractions 8 to 16 in all our samples (**Figure S4**). Based on this observation, fractions 1 to 7 corresponding to the non-polysomal contents and fractions 8 to 16 corresponding to polysomal contents were pooled. We then quantified the levels of the HBV SPs and *GAPDH* as positive control by ddPCR. As expected, *GAPDH* mRNAs were segregated in polysomal fractions (**Figure 3a**). Strikingly, we also detected signals for all the tested HBV SPs in the polysomal fraction in both control and DDX5/17 –silenced cells in the absence of EDTA. Similarly to our positive control, this signal appeared to be decreased in the EDTA-treated control concomitantly to an increased proportion in non-polysomal fractions, validating our experimental approach (**Figure 3a**). Finally, DDX5/17 silencing had no impact on the proportion of these SPs in the polysomal fraction showing that DDX5 and DDX17 did not regulate their translation (**Figure 3a**). These data thus indicate that SP02, SP03, SP15, SP16 and SP17 are engaged in active translation and potentially encode new viral proteins.

**Figure 3:**
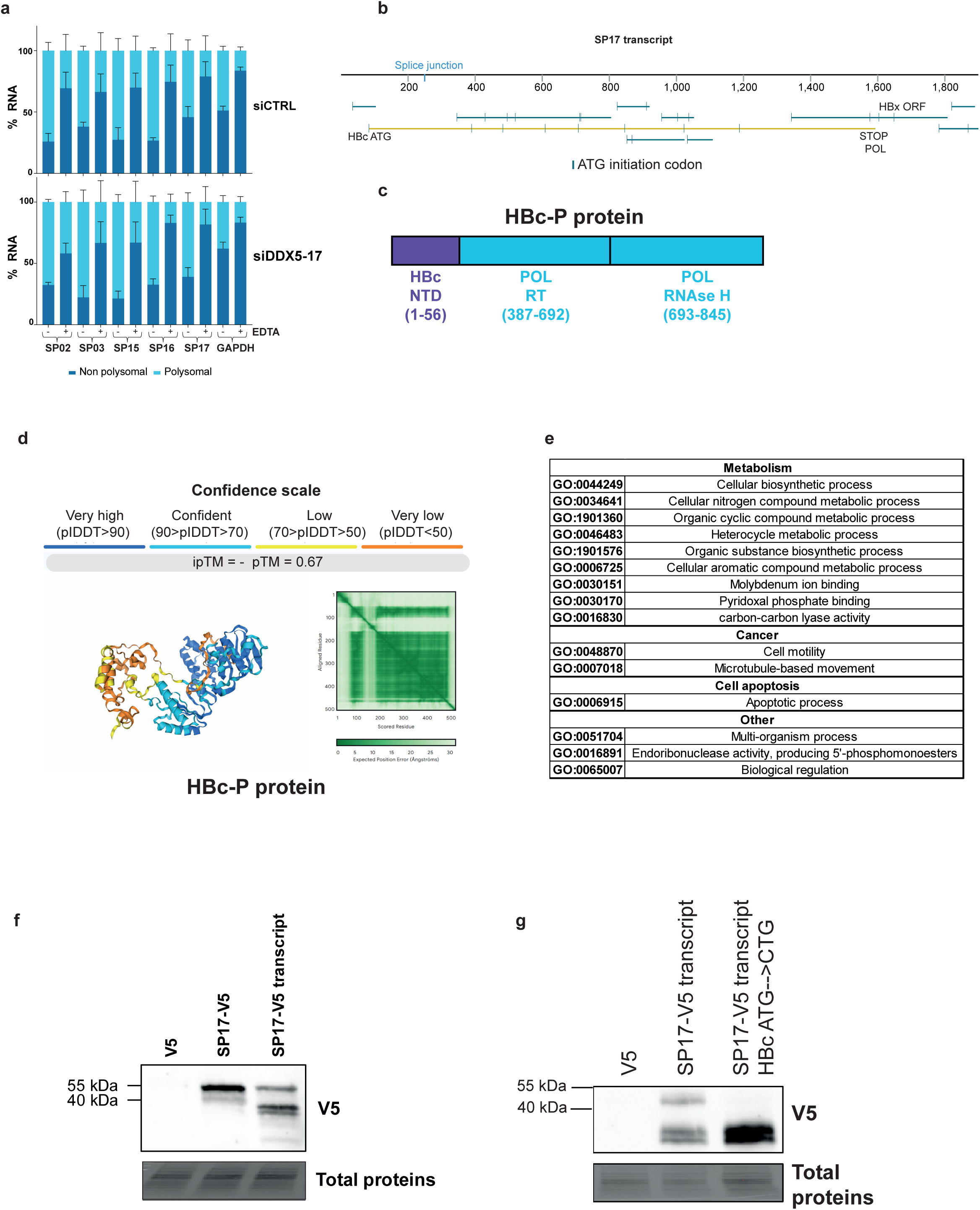
HBV SPs using the SD2070 as a donor site encode new viral proteins. **a** HepG2-NTCP cells treated as in Figure 1-3 were subjected to polysome fractionation. Polysomal (light blue) and non-polysomal (dark blue) fractions were pooled respectively and isolated RNAs were subjected to RT-ddPCR experiments to quantify SP02, SP03, SP15, SP16, and SP17 in siCTRL (top panel) and siDDX5-17 (bottom panel) conditions. Data are expressed as percentage of the Input. Error bars represent the standard error of the mean of two independent biological replicates. **b** The SP17 transcript contain several ORFs (green lines) including an ORF potentially encoding a fusion protein between HBc and POL, which we named HBc-P (yellow line). Vertical green lines correspond to ATG initiation codon **c** Protein domains of the HBc-P protein originated from the fusion between the N-terminal domain of HBc (HBc NTD), a truncated RT domain of POL (PLO RT) and the RNAse H domain of POL (POL RNAseH). Domain coordinates correspond to the coordinates of the parental proteins. **d** Tridimensional structure of HBc-P predicted from the HBc-P CDS using AlphaFold3 server (https://alphafoldserver.com/) ^38^. Above the structure is represented the confidence scale of this structure. **e** Biological functions of HBc-P protein predicted from the HBc-P CDS using ProteInfer ((https://google-research.github.io/proteinfer/) ^39^. **f** HepG2-NTCP cells were transfected with pcDNA6 plasmids expressing either the V5 tag alone (V5), the HBc-P CDS (SP17-V5) or the full SP17 transcript with a V5 tag in frame with the HBc ATG (SP17-V5 transcript). Two days after, proteins were extracted and subjected to a Western blot using the anti-V5 antibody. Total proteins were used as a loading control. **g** HepG2-NTCP cells were transfected with pcDNA6 plasmids expressing either the V5 tag alone (V5), or the full SP17 transcript with a V5 tag in frame with the HBc ATG (SP17-V5 transcript) carrying a wt (SP17-V5 transcript) or a mutant HBc ATG (SP17-V5 transcript HBc ATG→CTG). Two days after, proteins were extracted and subjected to a Western blot using the anti-V5 antibody. Total proteins were used as a loading control.

We then looked at the open reading frames (ORFs) contained in these particular SPs. Each SP contains multiple ORFs that start at different ATGs and with different lengths. Like every HBV transcript, every SP contain the HBx ORFs (**Figure 3b, S5**). As expected from all variants bearing the splicing junction between the DS2449 and the acceptor site at the position 489, SP02 and SP15 potentially encode the HpZ protein that was previously described as a HBV self-restriction factor ^11^ (**Figure S5**). In addition to this protein, SP02 and SP15 also contain an ORF potentially encoding a truncated HBc protein deleted from its last cysteine residue (HBc^-Cys^) that was recently shown to be part of the viral capsid and to facilitate the release of the HBV genome ^9^ (**Figure S5**). In addition to this deletion, the SP02 HBc ORF also carries a deletion between the aminoacids 57-150 corresponding to a region containing part of the N-terminal domain, the linker domain and part of the C-terminal domain (**Figure S5**). Similarly, the HBc ORFs contained in the SP15 also bears a deletion between aa 57 and 113 corresponding to part of the N-terminal domain (**Figure S5**). SP03, SP16 and SP17 contain ORFs encoding different proteins containing a truncated RT domain and a full RNAse H domain of the HBV POL protein fused at its N-terminal to the 56 first aa of HBc corresponding to the N-terminal domain (SP03 and SP17) (**Figure 3b-c, S5**) or to 8 aa starting at an unprecedented described ATG (SP16) (**Figure S5**). Finally, SP16 also contains the ORFs potentially encoding the same truncated HBc protein as the SP02 (**Figure S5**).

We then intended to predict the tri-dimensional structure of the above-mentioned SP-derived protein by AlphaFold3 and to compare it to either the predicted full length HBc or the POL structure ^38^. Although some parts of the different proteins were predicted with less confidence compared to others, we could notice some similarities and differences between SP-derived proteins and the full-length HBc and POL proteins (**Figure 3d, S6a-g**). In particular, the structure corresponding to the RNAse H domain was highly similar between the SP-derived proteins carrying this domain and POL. This strongly suggests that these new SP-derived proteins could potentially retain a part of the function of the full-length proteins but also display substantial differences.

We thus aimed to predict the functions of the SPs-derived proteins according to their aa sequence using ProteInfer ^39^ that uses deep convolutional neural network to predict functions from unaligned sequences. As expected, SP02 and SP15-derived proteins shared common predicted functions related to gene expression and viral entry (**Figure S7**) suggesting that they could regulate both viral and host biology. Interestingly, SP03, SP16 and SP17-derived proteins, that carry a full RNAse H domain, had predicted metabolic functions important for hepatocyte homeostasis (**Figure 3e, S7**). Besides, SP03 and SP17-derived proteins also had predicted functions in cell cycle and motility, a key aspect for cancer development (**Figure 3e, S7**). Altogether, these data suggest that the SPs-derived proteins could play a role in viral replication and HBV-induced liver pathogenesis.

### SP17 encodes the HBc-P protein

We finally aimed to validate the production of new HBV proteins by these SPs, and particularly focused on the SP17 that is the strongest SP induced by DDX5/17 silencing in hepatocytes (**Figure 2b-c**). To this aim, we engineered two constructs carrying either the coding sequence of the SP17 protein we called HBc-P fused to a V5 moiety at its C-terminal end (HBc-P) or the full SP17 transcript carrying a V5 sequence in frame with the HBc ATG predicted to initiate the translation of HBc-P (SP17 transcript). We then transfected HepG2-NTCP cells with these constructs or with a control plasmid expressing only the V5 tag. Two days later, we performed Western blot analysis using an anti-V5 antibody. Interestingly, a single band at the expected size of HBc-P was detected in cells transfected with the HBc-P construct but not the in the control plasmid. This band was also detected in cells transfected with the SP17 transcript construct alongside with two smaller extra bands at around 40 kDa (**Figure 3f**). To validate the production of HBc-P protein by the SP17 transcript, we engineered another construct carrying mutations in the HBc ATG initiation codon and transfected HepG2-NTCP cells with this construct and the wt counterpart. Western blot analyses using an anti-V5 antibody revealed that, while the band corresponding to HBc-P is only present in cells transfected with the wt construct, the two other bands are present in both conditions (**Figure 3g**). These data thus confirmed the production of the HBc-P protein by the SP17. Furthermore, at least two other proteins starting at ATG in frame with the V5 tag are produced from this SP. According to their size, these proteins might correspond to additional proteins containing a truncated RT domain and the RNAse H domains. Taken together, these data demonstrated that SP17 encodes new HBV proteins potentially involved in the HBV life cycle and in cellular processes.

## DISCUSSION

Alternative RNA splicing is a potent mechanism that allows cells to adapt to their microenvironment by expanding their transcriptome and proteome ^40^. This process also appears to be essential for the replication of many viruses, such as HIV-1, that were described to hijack the splicing machinery ^41^. Albeit not essential for its replication, HBV also generates several spliced variants that were described to contribute to liver disease ^5,10^. Nonetheless, due to technological barriers, how this process is regulated in infected hepatocytes and which molecular mechanisms are involved in its contribution to HBV biology remain elusive. Thanks to our recently developed method based on HBV-specific 5’RACE-PCR assays that we combined to ONT-mediated single molecule sequencing, we were able to estimate the proportion of each individual HBV SPs in a relevant infection model in order to interrogate the contribution of specific host factors in the regulation of HBV RNA splicing ^4,34^.

We focused on DDX5 and DDX17 RNA helicases that we and other groups previously identified as HBV restriction factors ^31–33^. In particular, we recently demonstrated that both helicases are critical regulators of HBV RNA metabolism and, more precisely, of the processing of their 3’ end extremity ^31^. We thus investigated the role of both proteins in HBV RNA splicing by using our approach in HepG2-NTCP cells. More precisely, we demonstrated that the recognition of a specific splicing donor site at the position 2070 is strengthened upon DDX5/17 silencing. This site is engaged in splicing junction present in SP02, SP03, SP08, SP10, SP12, SP15, SP16 and SP17. Accordingly, except for SP08 and SP12 that were not detected in HepG2-NTCP cells, silencing of both helicases resulted in a general increase of these SPs, SP17 showing the highest induction (from < 1% to 10-15 % of SPs). These data were validated by RNA-seq experiments and RT-ddPCR using specific junction primers in HepG2-NTCP cells. We also demonstrated that this peculiar role of DDX5 and DDX17 was conserved in PHH, a more physiological model of HBV infection. This observation raises the obvious question about the mechanisms that might confer this specificity of action of both helicases on this DS2070. Recently, silencing of both DDX5 and DDX17 was also associated with defects in alternative splicing and 3’ end mRNA processing on a subset of mammalian genes in neuroblastoma cells. This was functionally linked to the role of both helicases in the tridimensional chromatin organisation of the regulated loci pointing out the role of DDX5 and DDX17 in coupling chromatin and co-transcriptional RNA processing ^42^. Interestingly, the DS2070 is located 150 bp downstream of the poly-A signal whose recognition by termination factors is impeded by DDX5 and DDX17 ^31^. These observations thus suggest that both helicases, bound to cccDNA, regulate the topology of this particular region to mask important regulatory signals that dictate the fate of HBV RNAs. Testing this hypothesis is still technologically challenging due to the small size of the HBV minichromosome. This proposed mechanism does not exclude a more direct role of DDX5 and DDX17 on HBV RNAs. Indeed, we and another group previously demonstrated that DDX5 and DDX17 are also bound to HBV RNAs in HepG2-NTCP cells ^31,32^. These proteins are RNA helicases whose main functions are to resolve secondary structures^22^. The resolution of such structures facilitates the passage of the RNAP II enhancing its elongation speed and preventing the recognition of weak splicing sites by the spliceosome ^43^. Interestingly, DDX17 was found to bind to the ε loop of the pgRNA, an essential structure for HBV biology located around 200 bp upstream of the DS2070 (position 1849-1909) ^32^. Thus, by resolving the pgRNA ε loop, DDX17 and, potentially, DDX5, which is functionally redundant with DDX17 ^23,31^, would increase the RNAP II elongation rate at this particular region and prevent the recognition of the DS2070 in infected hepatocytes.

DDX5 and DDX17 appeared to be down-regulated in CHB patients and low DDX5 expression in HBV-induced HCC patients is associated with poor prognosis ^31,33^. Based on our results, this would correlate with the increased production of the DS2070 containing SPs. The landscape of HBV SPs evolves across the different stages of CHB ^5,12,13^, suggesting an active role of these variants in HBV-induced liver pathogenesis. Interestingly, *in vitro* co-culture assays revealed that the expression of SP02 and SP03, two of the SPs induced by DDX5 and DDX17 silencing, in hepatocytes led to the activation of hepatic stellate cells, a key event in the induction of fibrosis ^13^. HBV SPs are thought to act via their encoded proteins. For instance, SP01, the main SP, encodes three different viral proteins that not only regulate HBV replication (HPz and HBc^-Cys^) but also modulate the host innate immune response against HBV (HBSP) ^9,11,15^. Here, by performing polysome fractionation, we demonstrated that the SPs regulated by DDX5 and DDX17 are translationally active and potentially encode new viral proteins. This results in the production of new HBC^-Cys^ truncated proteins or fusion proteins containing a truncated POL RT domain and a full RNAse H domain. Their predicted structure using AlphaFold3 showed similarities and differences with their original viral proteins probably highlighting the conservation of certain functions and the acquisition of novel roles in the context of CHB. Accordingly, the prediction of their function using ProteInfer suggested functions associated with metabolism, cancer development, and virion assembly. We validated by transfection assay the production from SP17, the mostly induced variant upon DDX5 and DDX17 silencing, of a fusion protein between HBc and Pol, that we named HBc-P. As mentioned above, this fusion protein contains a full RNAse H domain that was demonstrated to inhibit the response to interferon alpha opening the possibility that SP17 could also participate to lower the sensitivity of infected hepatocyte to this drug ^44,45^. Furthermore, prediction of HBc-P functions by ProteInfer suggested a potential role of this protein in metabolism and cancer and, thus, during liver pathogenesis. Further investigations will be required to determine the precise role of SP17 and the other SPs regulated by DDX5 and DDX17 in hepatocytes and in patients.

To conclude, we identified DDX5 and DDX17 as new regulators of HBV RNA splicing. Their down-regulation observed in CHB patients could be an essential step for the progression of the disease by allowing the expression of HBV SPs encoding new viral proteins actively contributing to HBV-induced liver pathogenesis.

## MATERIALS & METHODS

### Production of HBV viral inoculum

HBV (Genotype D, subtype ayw) inoculum was prepared from filtered HepAD38 supernatants by precipitation with 8 % polyethylene-glycol-MW-8000 (PEG8000, SIGMA) as previously described ^46^. Viral stock with a titer of at least 1×10^10^ viral genome equivalents (vge)/mL was tested endotoxin-free and used for infection.

### PHH isolation

Primary human hepatocytes were isolated from surgical liver resections after informed consent of patient (IRB agreements #DC-2008-99 and DC-2008-101) as previously described ^47^ and plated in complete William’s supplemented with 1 % penicillin/streptomycin (Life Technologies), 1% L-Glutamine (Life Technologies), 5 µg/mL insulin (Sigma-Aldrich), 25 µg/mL hydrocortisone hemisuccinate UPJOHN (SERB) and 5 % fetal calf serum (FCS. Fetalclone II^TM^, PERBIO). PHH were maintained in William’s medium supplemented with 1.8% DMSO (Sigma-Aldrich). All PHH-related data were obtained from at least three distinct donors. All experiments on PHH were performed as described for HepG2-NTCP cells.

### Cell culture, HBV infection, siRNA transfection

HepG2-NTCP cells were cultured in proliferation medium (DMEM medium supplemented with 1 % sodium pyruvate (Life technologies), 1% Glutamax (Life technologies) and 5 % FCS (Fetalclone II^TM^, PERBIO)) at 37 °C and 5 % CO2. Three days prior to HBV infection, cells were seeded at 1.5 x 10^5^ cells/cm² in proliferation medium. The day after, medium was changed for differentiation medium (proliferation medium supplemented with 2.5 % DMSO). 48 h after, cells were infected with 250 vge/cell in presence of 4 % PEG8000 for up to 16 h. After three washes with 1 X PBS, cells were maintained in differentiation medium. Four days post-infection, cells were trypsinated, reseeded at 1.5 x 10^5^ cells/cm² in proliferation medium and transfected with 25 nM control siRNA targeting the luciferase mRNA (Horizon) or 12.5 nM siRNAs targeting DDX5 mRNA (Horizon # 4392420) and 12.5 nM siRNAs targeting DDX17 (Horizon # 4390824) mRNA using Lipofectamine RNAi Max (Life technologies) according to manufacturer’s instructions. The day after, cells were washed once with 1 X PBS and maintained in differentiation medium. Transfection was repeated the day after while cells were maintained in proliferation medium for an extra day. Then, cells were again washed once with 1 X PBS and cultured for a final day with differentiation medium.

### Plasmid and transfection

Plasmid carrying either the SP17 CDS alone fused to a V5 tag at its 3’ end extremity or the full SP17 transcript with a V5 tag in frame with the HBc ATG or the V5 tag alone were obtained by cloning the corresponding DNA fragment obtained from GenScript between the AgeI and HindIII restriction site of the pcDNA6/GFP plasmid previously described ^48^. HepG2-NTCP cells were reverse transfected with the above plasmids using the TransIT-2020 (Mirus) according to manufacturer’s instruction and cultured for two days in proliferation medium.

### RNA extraction

Cells were directly lysed in 1 mL RNAZol (MRC Reagent) for up to 10^7^ cells and 0.4 volume of H2O was added to the lysate. The mixture was then vortexed for 15 seconds, incubated for 15 minutes in ice and centrifuged 15 minutes at full speed at 4 °C. 1 mL was collected and RNA was precipitated with 0.4 volume of 75 % EtOH. After 15 minutes in ice, samples were again centrifuged 30 minutes at full speed at 4 °C. RNA pellet was washed twice with 70 % EtOH, centrifuged 5 minutes at full speed at 4 °C and air-dried for 5 minutes. RNA was finally resuspended in RNAse-free H2O and its concentration was estimated by Nanodrop One (Thermo Fisher).

### 5’ RACE-PCR

5’RACE-PCR experiments were performed following our previously described protocol ^49^. Primer sequences are listed in **Table 1**.

### Nanopore sequencing

5’RACE-PCR amplicons were prepared for Oxford nanopore sequencing (ONT) using the ligation sequencing gDNA kit (SQK-LSK109, ONT) and barcoded by PCR using the PCR Barcoding Expansion 1-12 kit (EXP-PBC001, ONT) according to the manufacturer’s instructions. Equimolar amount of each sample was then pooled together and the library was loaded on a MinION Flow Cell (MIN-106D R9, ONT) and sequenced for 72 h.

### Sequencing analysis

Nanopore sequencing data were analysed using the Bolero pipeline previously described ^34^.

### RNA-seq experiments and bioinformatics analysis

Total RNA isolated from HepG2-NTCP cells treated as above were depleted from ribosomal RNAs using the Illumina® Stranded Total RNA Prep, Ligation with Ribo-Zero Plus kit (Illumina) and the library was further prepared using NovaSeq XP 2-Lane Kit v1.5 (Illumina) according to manufacturer’s instructions. Samples were sequenced on a Nova Seq machine (Illumina). Reads aligned to HBV genome containing the different HBV RNA donor sites engaged in splicing junctions were counted and normalised to the number of reads aligned to a region that is specific to the 3.5 kb RNAs ^31^.

### Reverse transcription and ddPCR

1 µg RNA was treated with RQ1 DNAse (Promega) in 1 X RQ1 buffer (Promega) for 30 minutes at 37 °C. RQ1 was then inactivated by adding 1 X Stop solution and an incubation of 10 minutes at 65 °C. DNAse-treated RNAs were then reverse-transcribed using the SSIII 1ST strand qPCR supermix (Life technologies) according to the manufacturer’s instructions.

50 ng cDNA with 200 nM primers specific to HBV SPs (**Figure S4a**) or to 3.5 kb RNAs and 1X QX200^TM^ ddPCR^TM^ EvaGreen Supermix (Bio-Rad) were partitioned into the Automated Droplet Generator with specific oil for EvaGreen (Biorad), amplified in the C1000 Touch thermal cycler (Bio-Rad) following the program: 5 minutes at 95 °C, 40 cycles of 30 seconds at 95 °C and 1 minute at 60 °C, 5 minutes at 4 °C and 5 minutes at 90 °C. Analysis was then performed with the QX100 Droplet Reader (Bio-Rad) using the Quantasoft software (v1.7.4, Bio-Rad, Hercules, CA, USA). Absolute copy numbers of HBV SPs were then normalised with the absolute copy number of the 3.5 kb RNAs. Primer sequences are listed in **Table 1**.

### Polysome fractionation

Polysome fractionation was performed as previously described ^37^. Briefly, cells were incubated for 3 minutes in 1 X PBS supplemented with 100 µg/mL cycloheximide (Merck) (PBS-cyclo) at room temperature and washed twice with PBS-cyclo. Cells were then scrapped and centrifuged at 1000 g for 5 minutes at 4 °C. Cell pellet was then lysed in PLB lysis buffer by pipetting up and down twenty times. 50 mM EDTA was added in the negative controls and samples were incubated for 10 minutes in ice. After a centrifuge of 5 minutes at 10000 g for 5 minutes, the supernatant was delicately loaded on a 15-40 % sucrose and ultracentrifuged at 38000 rpm for 2 h at 4 °C with maximum acceleration and no brake in a SW41 Ti rotor. 700 µL fractions were collected and 1 volume of acidic phenol/chloroform was added. After vigorous shaking, samples were centrifuged for 15 minutes at 12000 g at 4 °C. The top phase was then collected and RNAs were precipitated by adding 1 volume of isopropanol containing glycogen. The mixture was then incubated overnight at – 20 °C and centrifuged for 1 h at 14000 g at 4°C. RNA pellets were washed once with 75 % EtOH, air dried for 5 minutes and resuspended in RNAse-free H2O. A small aliquot was then loaded on agarose gel to define polysomal and on polysomal fractions. The remaining samples were then treated with DNase, reverse transcribed and subjected to ddPCR as previously described.

### Western blot

Cells were washed twice with 1 X PBS, centrifuged 5 minutes at 1500 rpm and resuspended in RIPA buffer supplemented with a cocktail of protease inhibitors (Complete EDTA-free, Roche). After an incubation of 30 minutes in ice and centrifugation for 30 minutes at 4 °C at full speed, the supernatant containing the proteins were dosed using a BCA assay (Life technologies) according to the manufacturer’s instructions. 20 µg of proteins in 1 X Laemmli buffer were denatured for 5 minutes at 95 °C, loaded on a 4-20% MP TGX Stain-Free Gel (Bio-Rad) and run in 1 X TG-SDS buffer (Euromedex). Gels were activated for 45 seconds on a Chemidoc MP (Bio-Rad) and proteins were transferred in a nitrocellulose membrane using a Transblot Turbo machine using the mixed molecular weight program (Bio-Rad). Signals for total proteins were captured using the ChemiDoc MP (Bio-Rad). Membranes were blocked for 30 minutes in 5 % Milk/0.1 % Tween20/1 X TBS at room temperature and incubated in primary antibodies prepared in the blocking buffer for 2 h at room temperature (**Table 1**). After three washes with 1 X TBS/0.1 % Tween20 for 5 minutes, membranes were incubated for 1 h at room temperature with HRP-conjugated secondary antibodies prepared in the blocking buffer (**Table 1**). After three washes with 1 X TBS/0.1 % Tween20 for 5 minutes, membranes were incubated for 3 minutes with the peroxidase substrate (Clarity, Bio-Rad) and the chemiluminescence signal was detected using the ChemiDoc MP machine (Bio-Rad).

### Protein structure prediction

Structures of the proteins predicted to be produced by HBV SPs were predicted using the AlphaFold3 server (https://alphafoldserver.com/) by interrogating the CDS sequence using the default settings ^38^.

### Protein function prediction

Functions of the proteins predicted to originate from the HBV SPs were predicted using ProteInfer (https://google-research.github.io/proteinfer/) by interrogating the CDS sequence using the default settings ^39^.

### Data analysis

Experiments have been performed at least four times (except where indicated) and graphical representations have been performed using GraphPad Prism 9.4.1. Mann-Whitney or Kruskal-Wallis statistical tests have been used to test the significance of the difference between the different comparisons. p<0.05 was used as a threshold to define the statistical significance.

## ACKNOWLEDGMENTS

We would like to thank Maud Michelet, Jennifer Molle, Anaëlle Dubois, Audrey Diederichs, Sarah Heintz and Mélanie Rodà for their help in the isolation of primary human hepatocytes, as well as Prof. M. Rivoire’s surgical staff for providing liver resections. The authors want to thank also all the members of the INSERM U1350 for the fruitful discussion. This study was supported by a public grant attributed by the French Agence Nationale de la Recherche (ANR) as part of the second “Investissements d’Avenir” program (reference: ANR-17-RHUS-0003) to FZ and by “Investissement d’avenir” Laboratoires d’Excellence (LabEx) DEVweCAN (Cancer Development and Targeted Therapies) grant ANR-10-LABX-61 to FZ and BT; by Agence Nationale de Recherches sur le SIDA, les Hépatites Virales et les Maladies Infectieuses Emergentes (ANRS MIE) grant ECTZ75178 to BT and CB and fellowship ECTZ161842 to GG.

## AUTHORSHIP CONTRIBUTION

Conceptualisation and formal analysis: GG, BT. Funding acquisition: GG, BT, CFB, FZ. Investigation: FC, GG, PH, MR, AD, RP, FDN. Methodology: GG, FC. Bioinformatics: XG. Resources: CS, IC, FZ, MR. Visualisation: GG, BT. Writing – original draft: GG, BT. Writing – review and editing: all authors.

## DATA AVAILABILITY STATEMENT

The data presented in this manuscript are available through the corresponding author upon reasonable request.

---

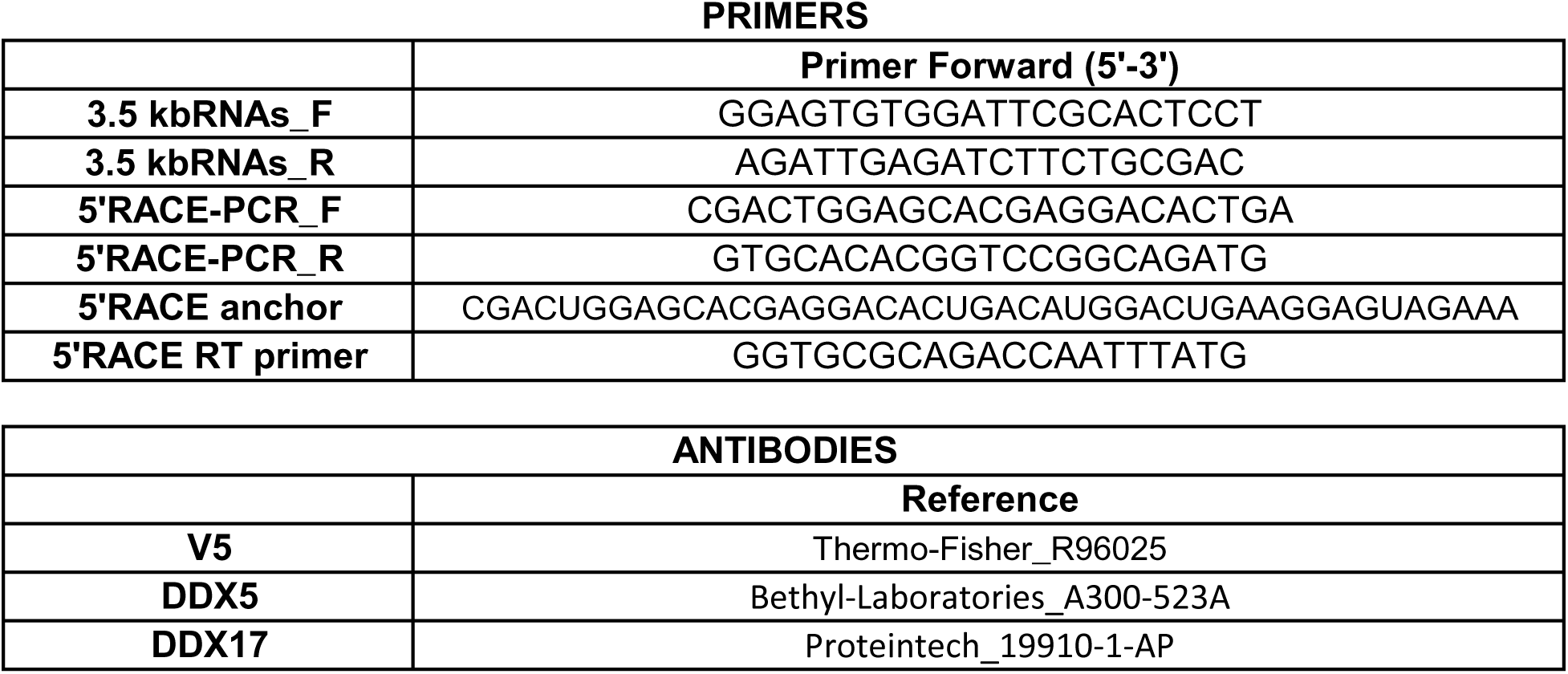

