## Supplementary material for "DDX5 and DDX17 RNA helicases regulate hepatitis B virus RNA splicing"

**SUPPLEMENTARY FIGURE LEGEND**

**Figure S1 (related to Figure 1): DDX5 and DDX17 silencing modifies the splicing landscape of HBV RNAs**

HepG2-NTCP cells were infected with HBV and transfected twice with control siRNAs (siCTRL) or siRNAs directed against DDX5 and DDX17 mRNAs (siDDX5-17).

**a** 8 days post-infection, proteins were extracted and subjected to Western blot analyses using an anti-DDX5 (top panel), an anti-ACTB (middle panel, loading control) or an anti-DDX17 (bottom panel) antibody. The figure is representative of five independent biological replicates.

**b** 8 days post-infection, RNAs were isolated and subjected to 5’ RACE-PCR. Amplicons were subjected to agarose gel electrophoresis to determine the profile of HBV transcripts. A 1 kb plus DNA ladder (Ladder) and a H2O-negative control were run in parallel. This panel is representative of five independent biological replicates.

**c** Percentage of each HBV SVs over their originated parental HBV transcripts in siCTRL (grey boxplot) and siDDX5-17 (green boxplot) conditions. Boxplots represent the minimum, the median and the maximum values of five independent biological replicates. Kruskal-Wallis tests with multiple comparisons were performed to test the difference between siCTRL and siDDX5-17 conditions. ns: p>0.05; **: p<0.01.

**d** Coverage of the TSS usage profile in siCTRL (top panel) and in siDDX5-17 (bottom panel) conditions.

**e** Percentage of the reads containing the different splicing donor sites engaged in a spliced junction in siCTRL and siDDX5-17 conditions. Kruskal-Wallis tests with multiple comparisons were performed to test the difference between siCTRL and siDDX5-17 conditions. ns: p>0.05; *: p<0.05.

**f** Percentage of the HBV reads containing the SD461 engaged in a splicing junction over the total number of preS2/S reads in siCTRL (grey boxplot) and siDDX5-17 conditions (green boxplot). Dashed lines connect each biological replicate.

**g** Percentage of SP21 and SP22 SPs containing the SD461 engaged in a splicing junction over the total number of preS2/S reads in siCTRL (grey boxplot) and siDDX5-17 conditions (green boxplot). Dashed lines connect each biological replicate.

**f-g** Boxplots represent the minimum, the median, and the maximum values of five independent biological replicates. Mann-Whitney were performed to compare siCTRL and siDDX5-17 conditions. ns: p>0.05; *: p<0.05; **: p<0.01.

**Figure S2 (related to Figure 2**): **DDX5 and DDX17 silencing leads to the differential usage of splicing donor sites of HBV RNAs**

GSE239571 RNA-seq dataset was reanalysed completed with two other biological replicates of HepG2-NTCP cells treated as in Figure S1. Reads containing the different HBV splicing donor sites engaged in a spliced junction were counted and normalised to the total number of reads aligning to HBV in siCTRL (grey boxplot) and siDDX5-17 (green boxplot) conditions. Kruskal-Wallis tests with multiple comparisons were performed to test the difference between

**Figure S3 (related to Figure 2**): **Validation of HBV SPs specific junction primers**

**a** Junction primers used to quantify specifically HBV SVs regulated by DDX5 and DDX17. The sequences in blue correspond to the sequence after the spliced junction.

**b** HepG2-NTCP cells were infected (HBV) or not (MOCK) with HBV for 8 days. RNAs were extracted and subjected to RT-ddPCR to amplify 3.5 kb RNAs, SP02, SP03, Sp15, SP16 and SP17 HBV SVs. A RT minus (RT-) and a H2O negative control were used in parallel to test the specificity of the amplification.

**Figure S4 (related to Figure 3): Polysome fractionation of control and DDX5-17-silenced HepG2-NTCP cells**

HepG2-NTCP cells treated as in Figure S1 were subjected to polysome fractionation. RNAs isolated from individual fractions were subjected to agarose gel electrophoresis to define the polysomal fractions (green).

**Figure S5: Predicted proteins produced from HBV SPs**

The HBV SVs regulated by DDX5 and DDX17 contain multiple ORFs (green line) including the HBx ORF and ORF encoding either truncated HBc proteins deleted of its last Cys residue (SP02 and SP15, orange line) or fusion proteins POL and HBc (SP03, orange line) or 8 aminoacid (SP16, orange line). Below the transcripts, are represented the different protein domains of the predicted proteins encoded by the orange ORFs. Coordinates numbers correspond to the coordinate of the parental protein.

**Figure S6: Predicted structure of the HBV SPs-derived proteins**

**a** Confidence scale of the predicted structure observed in **b-g**.

The tridimensional structures of the SP02 (**b**), SP03 (**c**), SP15 (**d**), SP16 (**e**), HBc (**f**) and POL (**g**) were predicted from the CDS of the corresponding proteins with AlphaFold3 (<https://alphafoldserver.com/>) (Abramson et al., 2024).

**Figure S7: Predicted biological functions of the HBV SPs-derived proteins**

Biological functions of the SP02, SP03, SP15 and SP16 proteins predicted from their CDS using ProteInfer ((<https://google-research.github.io/proteinfer/>) (Sanderson et al., 2023).
