## Supplementary figures and images for "DDX5 and DDX17 RNA helicases regulate hepatitis B virus RNA splicing"

### Figure S1

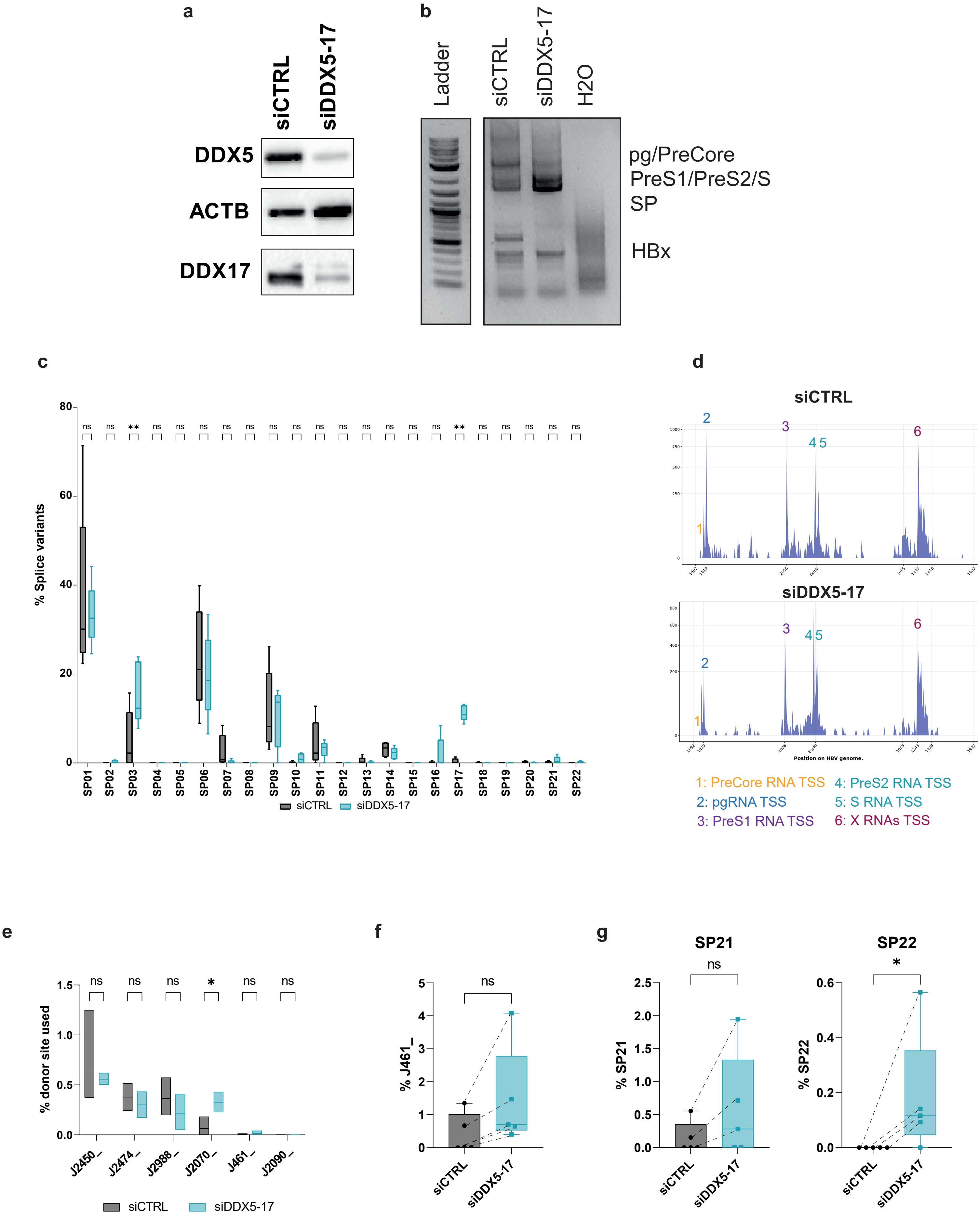

### Figure S2

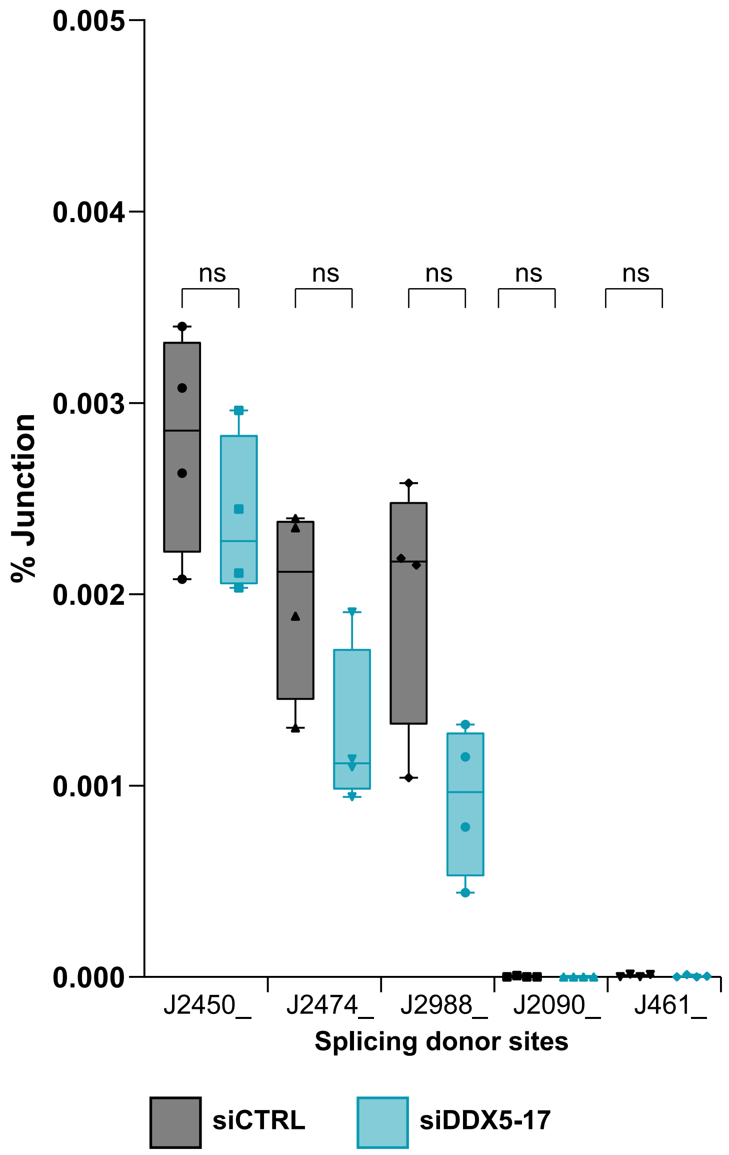

### Figure S4

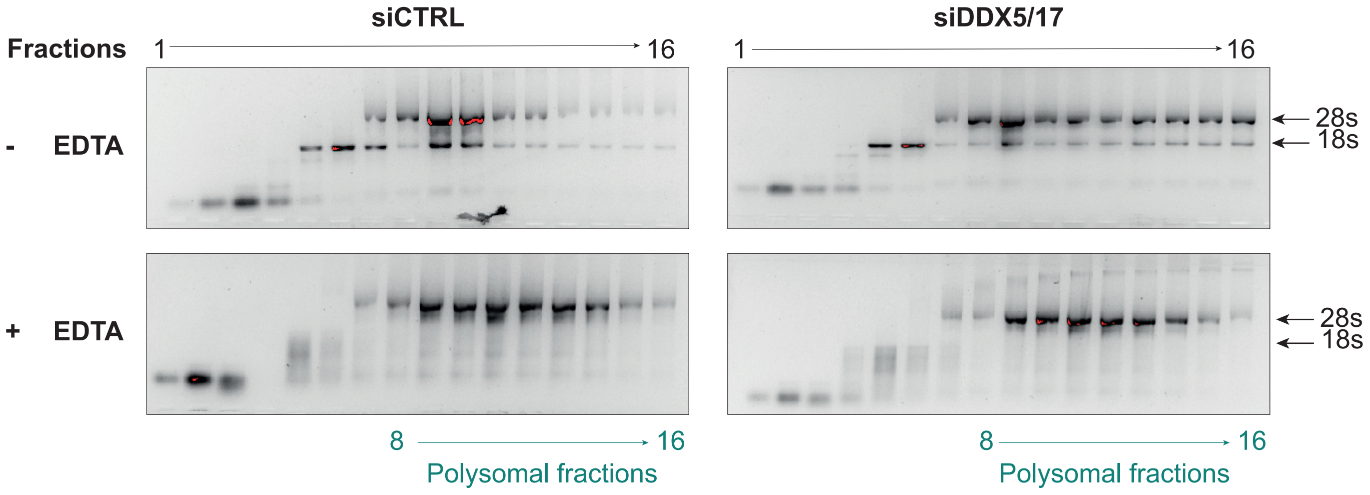

### Figure S5

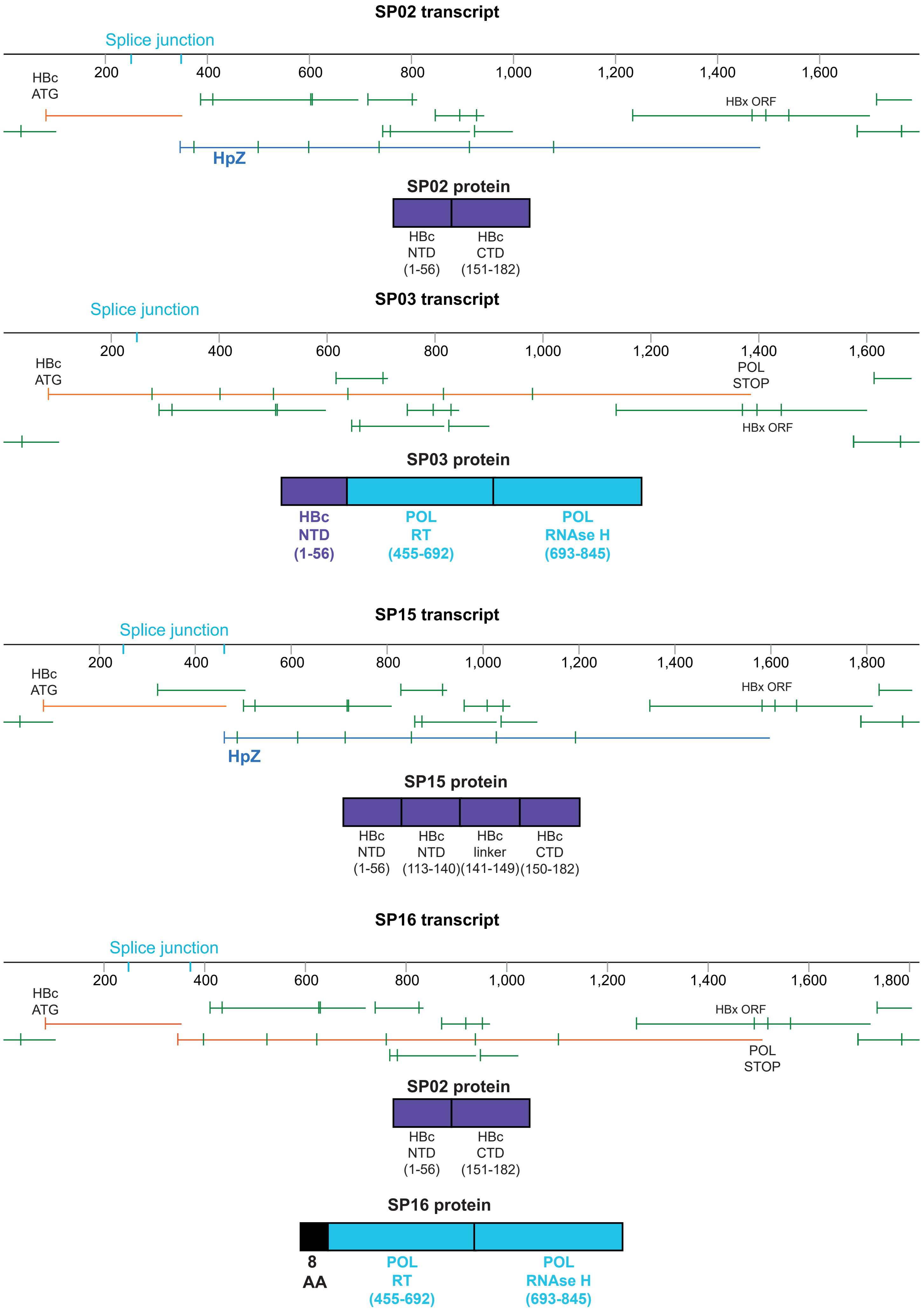

### Figure S6

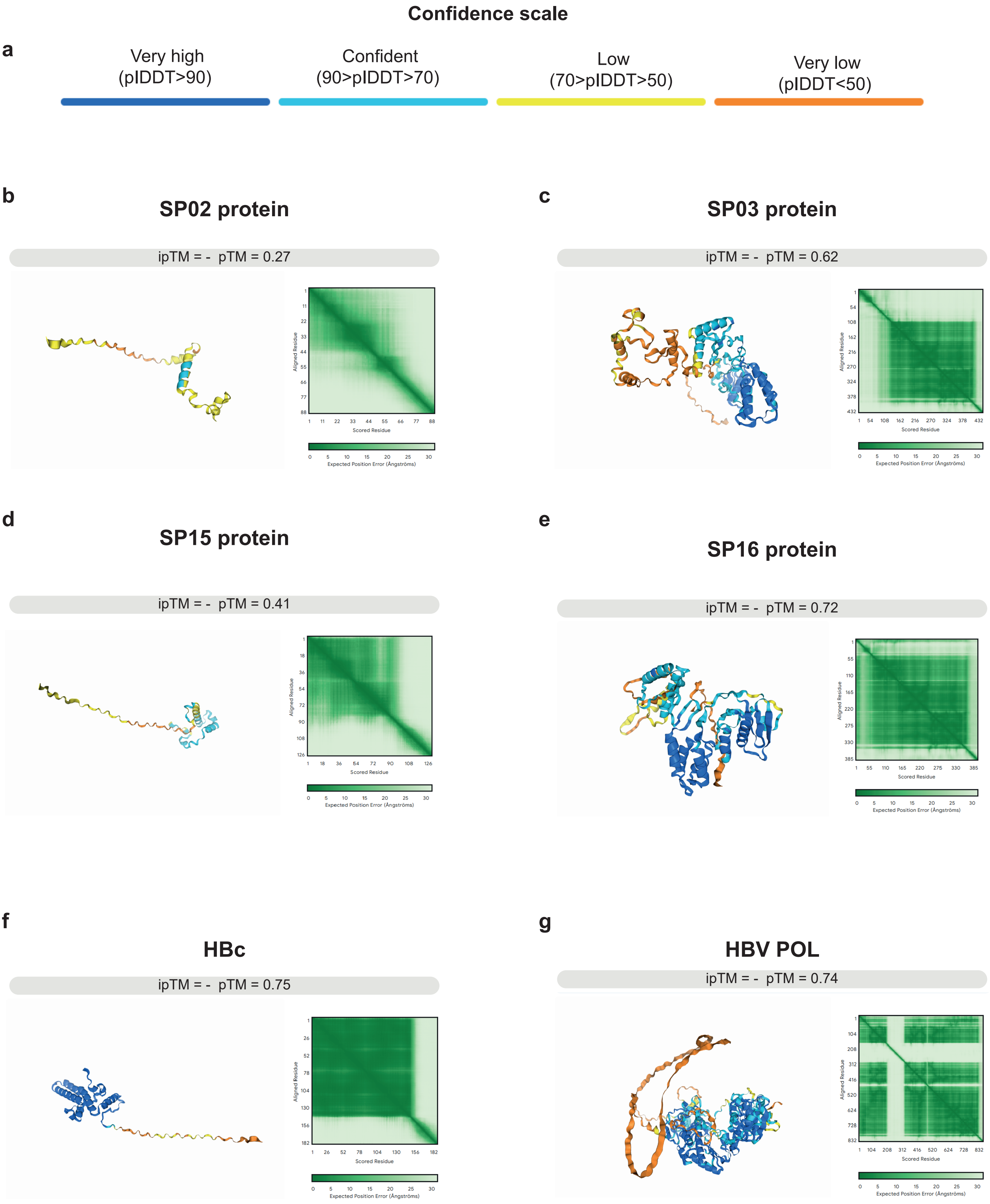
