## Supplementary material for "DDX5 and DDX17 RNA helicases regulate hepatitis B virus RNA splicing": Figure S3

a

| | Forward primer sequence | Reverse primer sequence |
|---|---|---|
| **SP02** | ACCATACTGCACTCAGAC**GACG** | GTGCTGGTGGTTGAGGAT**CATTGAG** |
| **SP03** | CACCATACTGCACTCAG**GATCCT** | GAGGCCCACTCCCATAGGAATT |
| **SP15** | CCTCACCATACTGCACTCAG**AGAAAC** | GTGCTGGTGGTTGAGGAT**CATTGAG** |
| **SP16** | ACCATACTGCACTCAGAC**GACG** | CTGGTGGTTGAGGAT**CCTTATG** |
| **SP17** | CACCATACTGCACTCAG**GGGGAA** | AGGACAAACGGGCAACATACCT |

b

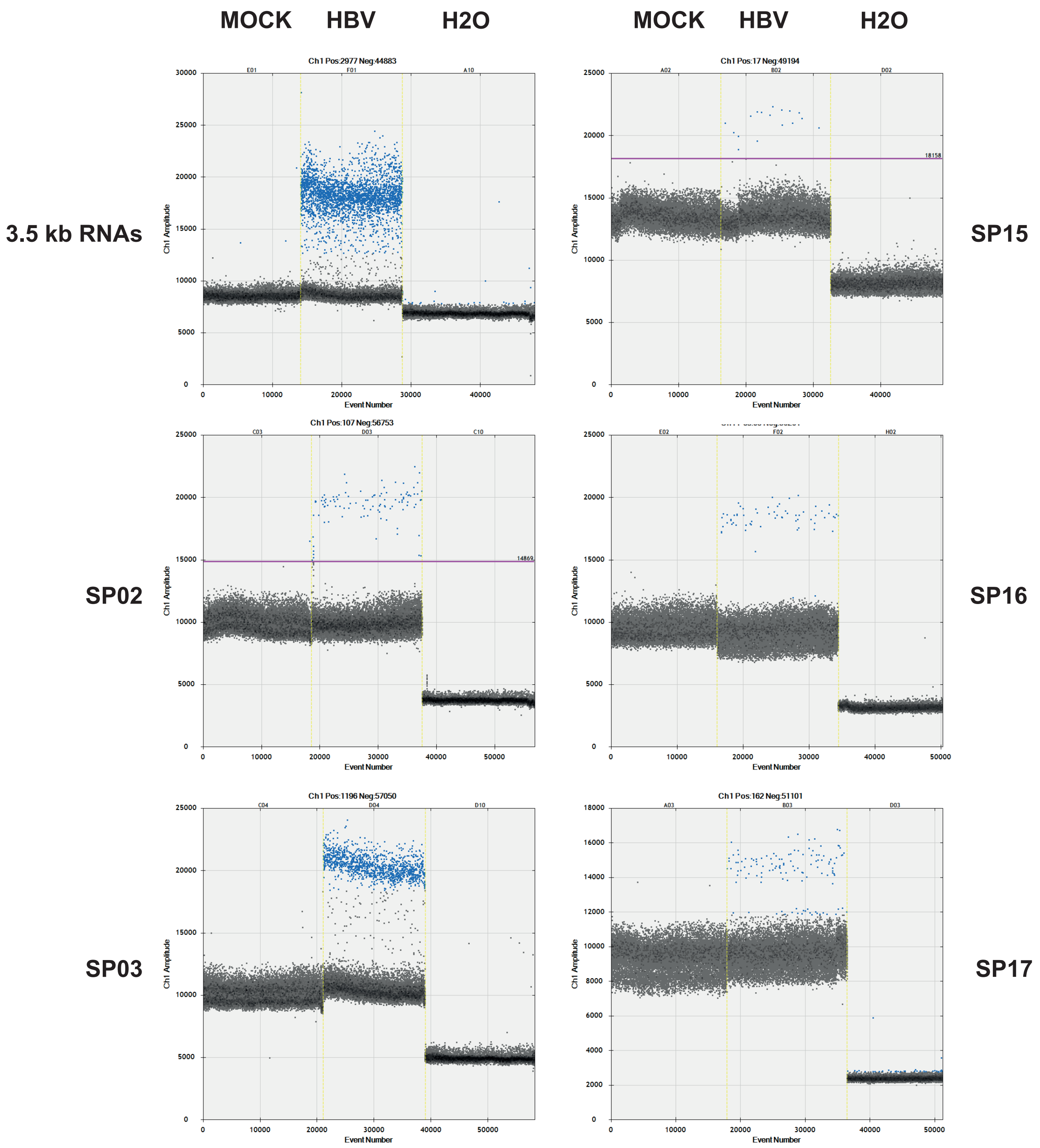


**Figure S3**
