## Supplementary material for "DDX5 and DDX17 RNA helicases regulate hepatitis B virus RNA splicing": Figure S7

| | |
|---|---|
| **SP02** | |
| **Apoptosis** | |
| GO:0043280 | Positive regulation of cysteine-type endopeptidase activity involved in execution phase of apoptosis |
| GO:2001270 | Regulation of cysteine-type endopeptidase activity involved in execution phase of apoptosis |
| **Gene expression** | |
| GO:0006323 | DNA packaging |
| GO:0003676 | Nucleic acid binding |
| GO:0016578 | Histone deubiquitination |
| GO:0005686 | U2 snRNP |
| GO:0000786 | Nucleosome |
| **Virus** | |
| GO:0075732 | Viral prenetration into host nucleus |
| GO:0019012 | Virion |
| GO:0044423 | Virion part |
| **Others** | |
| GO:0023014 | Signal transduction by protein phosphorylation |
| GO:0042025 | Host cell nucleus |
| GO:0033105 | Chlorosome envelope |
| **SP15** | |
| **Gene expression** | |
| GO:0016578 | Histone deubiquitination |
| GO:0000786 | Nucleosome |
| GO:0006337 | Nucleosome disassembly |
| GO:0034709 | Methylosome |
| GO:0031490 | Chromatin DNA binding |
| GO:0051276 | Chromosome organization |
| **Virus** | |
| GO:0039619 | T=4 icosahedral viral capsid |
| GO:0075732 | Viral penetration into host nucleus |
| GO:0075521 | Microtubule-dependent intracellular transport of viral material towards nucleus |
| GO:0019012 | Virion |
| **Others** | |
| GO:0090304 | Nucleic acid metabolic process |
| GO:0034622 | Cellular protein-containing complex assembly |
| GO:0021515 | Cell differentiation in spinal cord |
| GO:0071826 | Ribonucleoprotein complex subunit organization |
| GO:0030430 | Host cell cytoplasm |

| | |
|---|---|
| **SP03** | |
| **Cell cycle** | |
| GO:0007049 | Cell cycle |
| GO:0051301 | Cell division |
| GO:0051726 | Regulation of cell cycle |
| **Metabolism** | |
| GO:0034645 | Cellular macromolecule biosynthetic process |
| GO:0016830 | Carbon-carbon lyase activity |
| GO:0044255 | Cellular lipid metabolic process |
| GO:0008610 | Lipid biosynthetic process |
| GO:0009851 | Auxin biosynthetic process |
| GO:0030151 | Molybdenum ion binding |
| **DNA metabolism** | |
| GO:00140097 | Catalytic activity, acting on DNA |
| GO:0003677 | DNA binding |
| GO:0016779 | Nucleotidyltransferase activity |
| GO:2001251 | Negative regulation of chromosome organization |
| GO:0016891 | Endoribonulcease activity, producing 5'-phosphomonoesters |
| **Cell integrity** | |
| GO:0051494 | Negative regulation of cytoskeleton organization |
| GO:0044196 | Host cell nucleolus |
| GO:0000723 | Telomere maintenance |
| **Others** | |
| GO:0048070 | Regulation of developmental pigmentation |
| GO:0031362 | Anchored component of external side of plamsa membrane |
| GO:0048869 | Cellular developmental process |
| GO:0021915 | Neural tube development |
| **SP16** | |
| **Metabolism** | |
| GO:0034645 | Cellular macromolecule biosynthetic process |
| GO:001901363 | Heterocyclic compound binding |
| GO:0097159 | Organic cyclic compound binding |
| GO:0016830 | Carbon-carbon lyase activity |
| GO:0008299 | Isoprenoid biosynthetic process |
| **DNA metabolism** | |
| GO:0140097 | Catalytic activity, acting on DNA |
| GO:0051052 | Regulation of DNA metabolic process |
| **Others** | |
| GO:0031362 | Anchored component of external side of plasma membrane |
| GO:0000723 | Telomere maintenance |
| GO:0043167 | Ion binding |
| GO:0048869 | Cellular developmental process |
| GO:0007568 | Aging |
| GO:0051128 | Regulation of cellular component organization |

**Figure S7**
